# Different time scales of common-cause evidence shape multisensory integration, recalibration and motor adaptation

**DOI:** 10.1101/2023.01.27.525820

**Authors:** Nienke B Debats, Herbert Heuer, Christoph Kayser

## Abstract

Perception engages the processes of integration, recalibration and sometimes motor adaptation to deal with discrepant multisensory stimuli. These processes supposedly deal with sensory discrepancies on different time scales, with integration reducing immediate ones and recalibration and motor adaptation reflecting the cumulative influence of their recent history. Importantly, whether discrepant signals are bound during perception is guided by the brains’ inference of whether they originate from a common cause. When combined, these two notions lead to the hypothesis that the different time scales on which integration and recalibration (or motor adaptation) operate are associated with different time scales of evidence of a common cause underlying two signals. We tested this prediction in a well‐established visuo‐motor paradigm, in which human participants performed visually guided hand movements. The kinematic correlation between hand and cursor movements indicates their common origin, allowing us to manipulate the common‐cause evidence by this correlation between visual and proprioceptive signals. Specifically, we dissociated hand and cursor signals during individual movements while preserving their correlation across movement endpoints. Following our hypothesis, this manipulation reduced integration compared to a condition in which visual and proprioceptive signals were perfectly correlated. In contrast, recalibration and motor adaption were not affected. This supports the notion that multisensory integration and recalibration are guided by common‐cause evidence but deal with sensory discrepancies on different time scales: while integration is prompted by local common‐cause evidence and reduces immediate discrepancies instantaneously, recalibration and motor adaptation are prompted by global common‐cause evidence and reduce persistent discrepancies.

## Introduction

We frequently rely on multisensory signals when judging or manipulating objects. When these signals are discrepant our brain engages the processes of multisensory integration, sensory recalibration, and – when the signals relate to one’s own movements – also motor adaptation. Contemporary theories of multisensory perception propose that these processes are guided by the evidence about a common cause underlying both signals, as signals should be combined only when pertaining to a common object or action (Wallace et al., 2004; Kording et al., 2007; Chen and Spence, 2017). The body of multisensory research also proposes that these processes develop and operate on different time scales: while integration is thought to be essentially instantaneous and to reduce immediate discrepancies, recalibration and motor adaptation develop mostly over longer time scales and reflect the cumulative influence of recent multisensory discrepancies (Henriques and Cressman, 2012; Bosen et al., 2018; Rand and Heuer, 2019b; Noppeney, 2021; Debats et al., 2023). We here combine these two notions and hypothesize that the different time scales on which integration on the one and recalibration and motor adaptation on the other hand operate are associated with different time scales of evidence about a common cause.

The evidence about a common cause underlying two signals can have multiple origins, but prominently relies on spatio‐temporal correlations between two signals (Parise et al., 2012; Quintero et al., 2022). Indeed, artificially introduced correlations can enhance the binding of seemingly arbitrary signals (Ernst, 2007; Bizley et al., 2016; Tong et al., 2020; Kirsch and Kunde, 2022). Since the correlation of two signals can be manipulated on different time scales, this leads to the following prediction: when the correlation between multisensory signals is manipulated such that over shorter time scales the evidence for a common source is reduced, while over a longer time scale it remains intact, integration should be reduced but recalibration and motor adaptation should be preserved.

We here tested this prediction in a well‐established visuo‐motor paradigm that relies on visually guided hand movements similar to those performed when controlling a mouse cursor on a computer screen. Despite the hand and computer cursor moving in different spatial planes, both signals are perceptually combined and the presentation of visually rotated feedback engages the processes of multisensory integration (Ladwig et al., 2013; Rand and Heuer, 2013; Debats et al., 2017b; Kirsch and Kunde, 2019, 2022), multisensory recalibration(Rand and Heuer, 2019b), and motor adaptation (Krakauer et al., 2000; Bock et al., 2003; Debats et al., 2023). In fact, the properties of multisensory integration and recalibration seen in this visuo‐motor paradigm are analogous to those observed in purely perceptual paradigms such as the audio‐visual ventriloquism effect and aftereffect (Chen and Vroomen, 2013; Bruns, 2019; Park and Kayser, 2022). The likely reason for the perceptual combination of visual and proprioceptive signals in the visuo‐motor paradigm is that the spatially discrepant felt movements of the hand and seen movements of the cursor are temporally correlated in the same way as felt and seen movements of our hand. That is, the kinematic correlation between hand and cursor movements signals their common origin. Indeed, in a previous study we showed that manipulating the moment‐by‐moment correlation of hand and cursor trajectories can reduce multisensory integration (Debats et al., 2017a).

In the present study we leveraged this manipulation to dissociate multisensory integration from recalibration and motor adaptation. We introduced a local decorrelation of visual and proprioceptive signals during individual movements while keeping both signals correlated across series of movement endpoints. Following the above hypothesis, we expected this manipulation to reduce integration, which relies on causal evidence over short time scales, but to preserve the strength of recalibration and motor adaptation, as these rely on causal evidence unfolding over longer time scales. We implemented this decorrelation by providing manipulated visual feedback about the movement trajectory, which included a constant rotation of the visual trajectory (required to measure multisensory biases) and featured an additional increasing and decreasing rotation that reached a maximum halfway through the movement and which was zero at the start and endpoint (Figure 1). With this variable visuo‐motor rotation the angular discrepancies between the positions of the hand and the cursor are dissociated instead of being constant during each movement, but the endpoints of hand movements and cursor motions are identical to those with a constant visuo‐motor rotation. Hence, on a moment‐by‐moment basis the two signals contain little evidence for a causal relation, while across trials the two sensory signals are strongly correlated.

**Figure 1.**
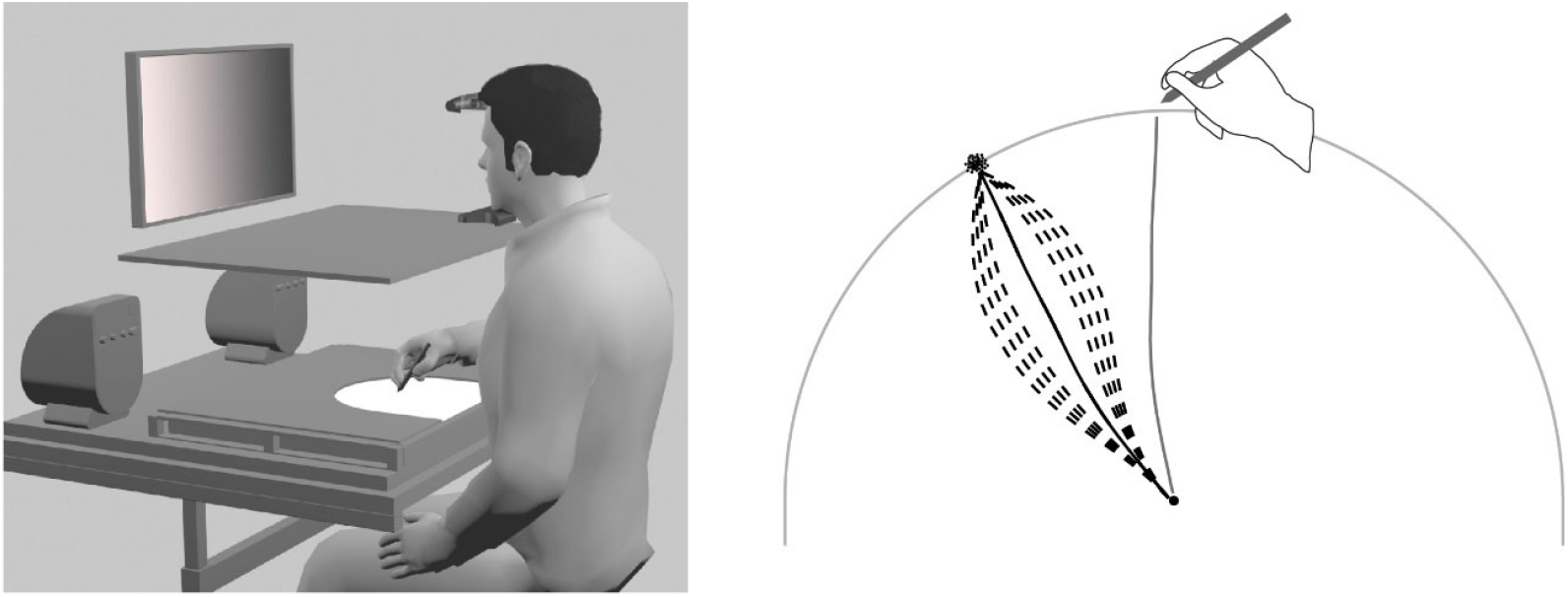
Apparatus and task. Left panel: The apparatus consisted of a semi‐circular workspace on which participants performed movements, a monitor for the presentation of visual feedback, a board to prevent participants from seeing their hand, and a chinrest to ensure a stable head position. Right panel: Participants performed center‐out movements (solid grey line) with rotated feedback (here by 30°). The kinematic trajectory of the visual feedback (a cloud of dots; indicated in black) was either at a constant angular offset (perfectly correlated) to the actual hand movement (condition constant; solid black line) or the trajectory was manipulated (condition variable; dashed black lines) such that the start and end points matched with the actual trajectory but the middle portion of the movement randomly deviated to the left or right (the example shows a number of variable trajectories for the one given movement).

## Methods

### Participants

We obtained data from 28 participants, aged 20 to 56 years (median: 26 years; 18 female). We did not plan this sample size a priori, as the expected effect sizes for recalibration and motor adaptation were not known. For integration our previous study suggests an effect size of about d=1.7 (Debats et al., 2017a), but those for recalibration and adaption may be smaller as these biases are generally weaker. According to G*Power 3.1.9.2 (Faul et al., 2007), a sample size of 28 results in a power of a one‐sided t‐test of 0.99 for an effect size of d=0.8 (at α=.05). Hence we deemed the present study sufficiently powered to replicate the previous result for integration and to detect potentially weaker effects for recalibration and motor adaptation. The experiments were conducted in accordance with the declaration of Helsinki and were approved by the Bielefeld University Ethics Committee. All participants reported being right‐handed, gave written informed consent prior to participation, and were compensated with a payment of €7 per hour.

### Apparatus

The apparatus was the same as used in previous studies using this paradigm (Debats et al., 2017a; Debats et al., 2021, 2023). Participants sat in front of a digitizer tablet (Wacom Intuos4 XL; 48.8 by 30.5 cm) and faced a computer monitor (Samsung MD230; 23 inches; 50.9 by 28.6 cm) at 60 cm viewing distance (Fig. 1, left). Their head was stabilized by a chin rest and they held a stylus with their right hand. When required, they pressed a button on this to submit a response. The stylus was moved on the digitizer tablet within a semi‐circular workspace (15 cm radius) bounded by a PVC board, and the position of the stylus was recorded with 60 samples per second (spatial resolution: 0.01 mm). Direct vision of hand and stylus was prevented by a horizontal opaque board, and the room light was switched off during the experiment. Stimuli presented on the monitor were in light grey on a black background except when noted otherwise. Stimulus presentation and movement recording were controlled by a custom‐made MATLAB program using the Psychophysics Toolbox (Brainard, 1997).

### Visuo‐motor task

Participants made center‐out and return hand movements on the digitizer tablet. Center‐out movements started at the center of the semi‐circular workspace, which was marked by an outline circle on the monitor, and were mechanically stopped by the workspace boundary (Fig. 1, right). In each trial one of five possible 15°‐ranges of directions was cued to prevent stereotyped movements. These ranges were centered at directions of 60, 75, 90 (forward), 105, and 120°. Participants were instructed to move smoothly and to remember the position where the hand hit the boundary (termed movement endpoint). These endpoints reflect the critical events on which the participant’s judgements and the analyses of interest are focused. There were no constraints on movement velocity. The return movements were from the boundary back to the remembered start position. Participants were instructed to return immediately upon hitting the boundary. This instruction served to prevent participants from spending variable and longer time intervals at the movement endpoints that could be used to memorize these. In some trials participants were asked to judge the endpoint of the center‐out movement of the hand or the cursor after the end of the return movement.

### Visual feedback and experimental conditions

Center‐out movements were performed with or without visual feedback presented on the monitor. In two experimental conditions different types of feedback were presented (Fig.1, right). In condition *constant* the direction of motion of the feedback was computed as D_c_(t) = D_h_(t)+C, with D_h_(t) as the direction of hand movement and C being a constant, either C=+30° or C=‐30°. This constant offset reflects the discrepancy between sensory signals that is required to engage multisensory processes. In this condition the correlation between the movement directions of hand and feedback marker was perfect. In condition *variable* the direction of motion of the feedback marker was D_c_(t) = D_h_(t)+C+A*sin(π*d(t)/d_max_), with A chosen randomly in each trial in the range of ‐15 to ‐7.5 or 7.5 to 15.0°. The proportion d(t)/d_max_ is the distance of the hand from the start position as a proportion of the maximal distance to the workspace boundary. At time point t=0 and when d(t)=d_max_ the term A*sin(π*d(t)/d_max_) is zero so that at the movement endpoints the visuo‐ motor rotations are identical in conditions *constant* and *variable*. However, during each movement in condition *variable* the kinematic correlation was reduced. The precise reduction was variable: it was stronger for larger amplitudes of the sinusoidal modulation of the rotation than for smaller amplitudes, and the relative reduction for positive and negative amplitudes depended on the curvature of the hand movements. The median correlation between the directions of hand and cursor movements was 0.735 (first and third quartile of the distribution: 0.445 and 0.894) in the experiment (computed across the 2800 exposure trials for condition *variable*). For both conditions the feedback marker was presented for 200 ms at the final position.

### Design

Each half of the experiment consisted of a pre‐test, an exposure phase, and a post‐test. The pre‐ and post‐test blocks contained bimodal trials with rotated visual feedback, used to probe multisensory integration, and unimodal trials, used to probe recalibration and motor adaptation. The exposure phase contained only bimodal trials, used to induce recalibration and motor adaptation. In the one half of the experiment, trials with visual feedback were of condition *constant*, and in the other half they were of condition *variable*. The order of these conditions was balanced across participants. For each sequence of the two conditions, two different sequences of constant rotations in the exposure phases were used, C=‐30° and C=+30°; thus, for each participant these constant rotations differed among the two experimental conditions.

The pre‐tests consisted of 40 trials with 8 permutations of the 5 cued ranges of movement directions. For the first half of the experiment, 20 bimodal trials were followed by 20 unimodal trials, for the second half this order was reversed. During bimodal trials visual feedback was presented with a rotation of C=+10 or C=‐10°, and for condition *variable* the within‐trial variation of the feedback rotation was added to the constant rotation; judgements of hand or cursor endpoints were requested (for each combination of C and type of judgement there were 5 trials with different cued ranges of movement directions). In 10 of the unimodal trials movements were performed without visual feedback, and in the other 10 trials only cursor motions were presented; participants were requested to judge the hand‐movement endpoint or the cursor‐motion endpoint, respectively.

The exposure phases consisted of two blocks of 50 trials each. In the first trials of the first block the constant rotation of the visual feedback both in conditions *constant* and *variable* increased in steps of +1° or ‐1° in each trial until +30° or ‐30° were reached. The gradual introduction of a visuo‐motor rotation is an established means to reduce or even prevent conscious awareness of the rotation and thus strategies to counter them (Kagerer et al., 1997; Buch et al., 2003; Michel et al., 2007). In 10 randomly chosen trials of each block participants were requested to judge the hand or cursor endpoint, in the remaining 40 trials no judgements were required. The judgements served to motivate participants to maintain attention to both the proprioceptive and visual endpoint information.

The post‐tests were identical to the pre‐tests with three exceptions. First, the order of bimodal and unimodal trials was reversed so that either the unimodal or the bimodal trials were adjacent to the exposure trials. Second, in bimodal trials the visuo‐motor rotations of +10° and ‐10° were shifted by one third of the constant rotation during exposure, that is, by +10° (+20 and 0°) or ‐10° (0 and ‐20°) to take the expected effects of the exposure phases into account. Third, after every 10 test trials five maintenance trials were inserted. These maintenance trials were identical to exposure trials (without judgements). The inclusion of bimodal trials in the post‐tests allowed us to re‐examine multisensory integration after a series of exposure trials which has been found to be essentially absent in a previous study (Rand and Heuer, 2020).

At the start of the experiment there was a block of 25 practice trials, for participants to familiarize with the task. Between the two halves of the experiment there was a block of 25 washout trials in which movements were performed with veridical visual feedback. No judgements were required. The total duration of the experiment was about two and a half‐hour. Pauses between trials were 2 s, and breaks between blocks were at the discretion of participants.

### Trial procedure

At the start of each trial an arrow guided the participant’s hand to an initial position by indicating the currently required direction of movement. The initial position was randomly selected within an area of 30*20 mm^2^, centered 30 mm nearer to the participant than the workspace center. After one second in the initial position, one of the five ranges of movement directions was cued. Cueing ranges of movement directions instead of clearly defined targets served to minimize the influence of targets on the perceived position of the hand. It should also minimize effects of actual movement errors (deviations of the visual‐feedback position from a target) on adaptive processes, but leave intact effects of prediction errors (deviations of the visual‐feedback position from its expected position) which are generally thought to be critical for adaptation (Shadmehr et al., 2010; Morehead and Orban de Xivry, 2021). The cue was a WiFi‐like symbol (three arcs of 15° widths at 18, 24, and 30 mm radial distance from the start position) which was presented for one second.

Following the presentation of the cue, an outline circle of 7 mm diameter was presented to mark the start position in the center of the workspace and a cursor (filled circle of 6 mm diameter) to indicate the participants’ hand position, enabling a visually guided movement from the initial position to the start position. This movement served to prevent participants from fixating a self‐selected target position for the center‐out movement. The colour of the filled circle was grey when the subsequent center‐out movement was accompanied by visual feedback, cyan when the center‐out movement was without visual feedback, and red in no‐movement trials in which only a center‐out motion of the feedback marker was presented. After 500 ms in the start position, the screen turned blank, cueing the participant to start the hand movement in movement trials.

The start of the center‐out movements was defined by leaving a tolerance range of 2.5 mm around the start position and their end by passing a threshold of 97% of the 150 mm distance between start position and workspace boundary. This threshold was chosen because the total movement distance varied slightly, depending on how the stylus was held. The center‐out movement was followed by a reversal phase that lasted from passing the 97% threshold in the outward direction until passing that threshold again in the backward direction. The return movement ended when the position of the hand did not change by more than 2.5 mm for 500 ms. Its end was signalled by a short beep.

During the exposure phase, the washout block, and in the bimodal trials of the test blocks and the practice block visual feedback was presented during the center‐out movements and for 200 ms in the final position. As described above, it was veridical or rotated by a constant or variable direction during each movement. The feedback marker had the format of a cloud of 100 grey dots (1 mm diameter each) the spatial coordinates of which had a bivariate normal distribution with standard deviations of 2 mm for both axes (truncated at three standard deviations so that the total diameter was about 12 mm). Visual feedback in the format of a cloud of small dots is only rarely used in visuo‐motor paradigms (Körding and Wolpert, 2004; Debats et al., 2021, 2023) but frequently in audio‐visual paradigms (Park et al., 2011; Rohe and Noppeney, 2015a; Beierholm et al., 2020; Park and Kayser, 2021, 2022) because it offers straightforward means to vary the reliability of the visual position information.

Judgements of the movement endpoints of hand or feedback marker were rendered in the pre‐ and post‐tests (and in a few trials of the exposure phases and the practice block). Following the return movement, the word ‘HAND’ or ‘CURSOR’ was presented on the monitor below the start position where it remained visible until the judgement was completed. One second after the word a grey filled circle of 2 mm diameter appeared at a random angular position on an invisible semi‐circle that corresponded to the workspace boundary. Participants adjusted the marker position along the semi‐ circular path by small movements of the hand‐held stylus relative to its position at the end of the return movement. The distance of the stylus from this position was proportional to the velocity of the marker (1 mm distance corresponded to a marker velocity of 1.5°/s). When the marker position was judged to match the remembered endpoint of the hand or feedback marker, respectively, participants pressed the stylus’ button to confirm their judgement. This was the end of the judgement provided the marker had been almost stationary (within a tolerance defined by a distance of 2.5 mm of the stylus from its position at the end of the return movement, corresponding to a velocity of 3.75°/s of the marker) for 500 ms. There were no time constraints for the judgements.

### Data pre‐processing

Data pre‐processing and analysis was similar to previous studies (e.g. (Debats et al., 2017a; Debats et al., 2021, 2023)). For each trial we recorded the movement trajectory and for judgement trials we recorded the participant’s endpoint judgements. All positions were transformed in polar coordinates centered on the start position of the center‐out movements. Except for data screening, only the angular positions were analysed because the distances of the physical and judged endpoints from the start position were constant throughout the experiment.

Ignoring the practice and washout blocks and the maintenance trials in the post‐tests, we collected a total of 4480 test trials and 5600 exposure trials. Half of them were in condition *constant* and the other half in condition *variable*. We screened these data for irregularities as follows: (1) the outward movement was not a smooth movement to the workspace boundary but included a reversal of more than 10 mm, (2) the stylus moved along the workspace boundary for more than 3.5° during reversal, (3) in trials in which only the feedback marker was presented the hand moved more than 15 mm, (4) the movement direction deviated more than 30° from the center of the cued range of movement directions, and (5) the absolute angular deviation between the physical and judged hand or cursor position was larger than 30°. The percentages of discarded trials were 3.4 and 3.1% for test trials in conditions *constant* and *variable*, respectively, and 4.3 and 3.7% for exposure trials.

### Analysis of multisensory integration

We analysed integration based on the angular judgement errors in bimodal pre‐test trials. Judgement errors were defined as deviations of the judged endpoints of the hand or feedback marker from the physical endpoints. We computed the integration bias as a proportion of the visuo‐motor rotation, defined as the difference between the angular judgement errors with visuo‐motor rotations of +10° and ‐10°, divided by the 20° difference between the physical rotations. If there is mutual attraction between the proprioceptive and the visual endpoint information, the proportional integration bias will be positive for the hand and negative for the visual feedback marker. Importantly, we expected the integration biases to be stronger in condition *constant* than in condition *variable* (Debats et al., 2017a).

To probe the influence of preceding exposure to discrepant signals on multisensory integration (Rand and Heuer, 2020), we also quantified the integration biases in the post‐tests. There, the perceived movements of the hand and the visual feedback should be affected by the preceding exposure phase with visuo‐motor rotations of +30 or ‐30°. In order to take sensory recalibration into account, the visuo‐ motor rotations in the bimodal trials of the post‐tests were shifted by +10 or ‐10°, with the effective rotations being 0 and 20° or ‐20 and 0°. From the findings of (Rand and Heuer, 2020) we expected that after an extended exposure period the integration biases should be reduced or even absent.

### Analysis of sensory recalibration

The analysis of proprioceptive and visual recalibration was based on the judgement errors in the unimodal trials of pre‐tests and post‐tests. For each participant the mean judgement error in the pre‐test was subtracted from the mean judgement error in the post‐test. For hand‐position judgements, these differences should be negative with a visuo‐motor rotation of ‐30° in the exposure phase and positive with a visuo‐motor rotation of +30°. For judgements of the feedback marker sensory recalibration should result in differences with the opposite sign. For the analysis of the full sample, the differences were multiplied with ‐1 when the exposure was with ‐30° rotation. Thus, for proprioceptive recalibration the sign of the differences should be positive, and for visual recalibration negative. If sensory recalibration was not affected by exposure to degraded kinematic correlations, the absolute recalibration biases would be similar in conditions *variable* and *constant*.

### Analysis of motor adaptation

The analysis of motor adaptation was based on the angular deviations of the movement endpoints from the centres of the cued ranges of movement directions (endpoint deviations) in the unimodal movement trials of pre‐tests and post‐tests. Differences between endpoint deviations between post‐test and pre‐test were analysed in the same way as for the analysis of sensory recalibration, but motor adaptation would be reflected in differences with the opposite arithmetic signs: a visuo‐motor rotation of ‐30° in the exposure phase should result in positive and of +30° in negative differences. After multiplication with ‐1 after exposure to ‐30°, the overall motor adaptation should be negative (it is also known as negative aftereffect).

### Statistical analyses

Statistical analyses were performed with custom‐written MATLAB code, STATISTICA 7.1, and JASP 0.16.4. Frequentist statistics were Student t‐tests for repeated measurements, effect sizes are quantified using Cohen’s d. These tests were one‐sided under the hypotheses that integration, recalibration, and motor adaptation were attenuated (or not) in condition *variable* as compared with condition *constant*. We report the t‐statistic with a positive sign when it is in the direction of the one‐sided alternative statistical hypothesis and with a negative sign otherwise. Mean values are reported with the Student’s t 95% confidence intervals in text (square brackets) and figures (error bars). Bayes factors B_0_ or B_1_ were computed for major findings that favoured the null or alternative hypothesis, respectively (using the default prior of JASP 0.16.4). When interpreting BF’s we refer to the nomenclature of (Wagenmakers et al., 2018): we interpreted BF between 1 and 3 as ‘anecdotal’, between 3 and 10 as ‘moderate’, and between 10 and 30 as ‘strong’ evidence. For the main comparisons between conditions *constant* and *variable* we added ANOVAs with order of conditions as additional factor to rule out that the differences between conditions were modulated by their order of presentation.

## Results

### Multisensory integration

We quantified integration based on judgements in the bimodal trials of the pre‐tests (Figure 2). Participants either judged the endpoint of the center‐out movements (‘hand judgement’) or the position of the visual feedback marker (‘visual judgement’). The errors for hand judgements were negative for negative (clockwise) visuo‐motor rotations and positive for positive rotations. Those for visual judgements were small and opposite to hand‐judgement errors, as expected.

**Figure 2.**
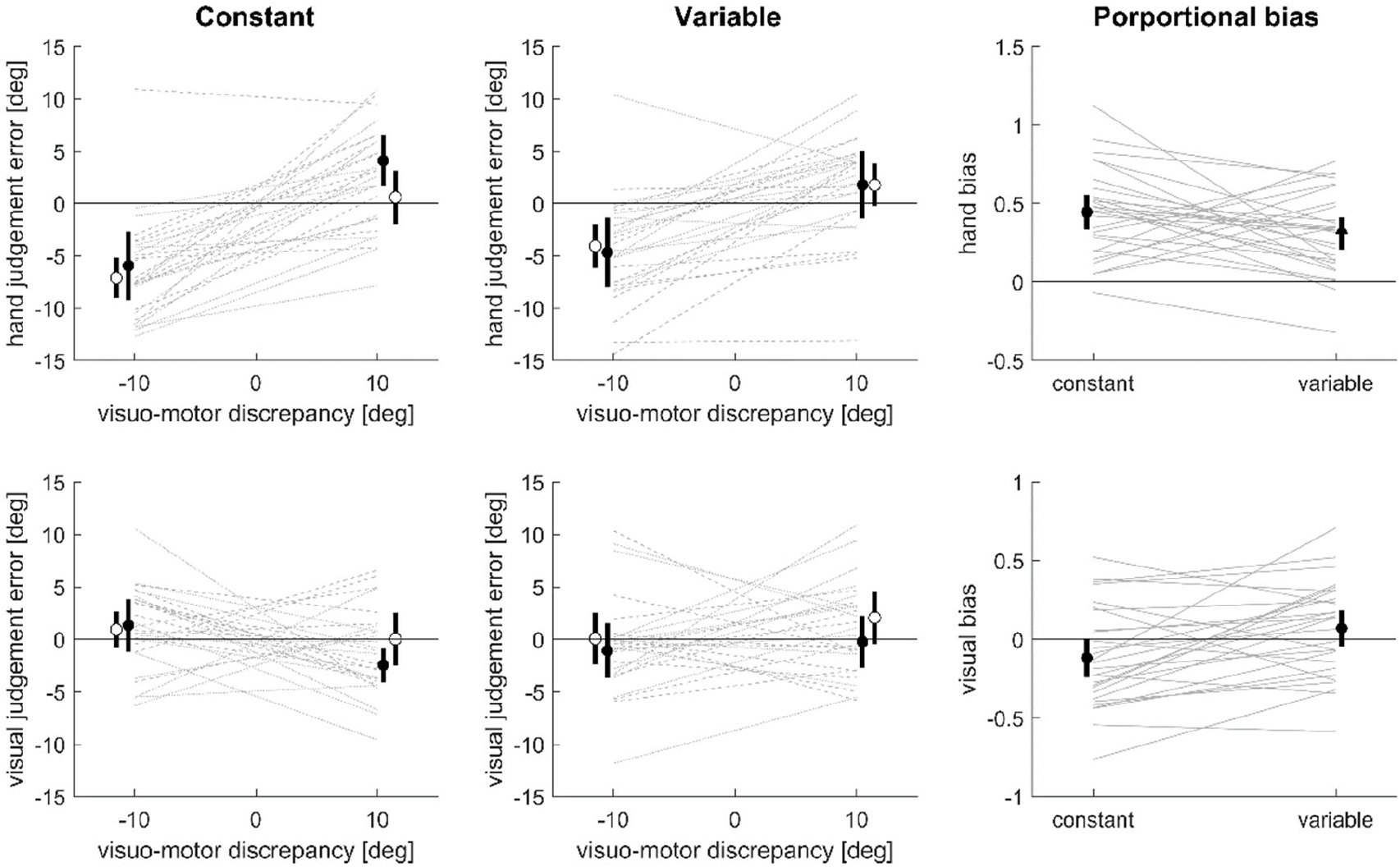
Multisensory integration. Left and middle panels: Judgement errors in bimodal pre‐test trials as a function of the visuo‐motor rotation for hand (upper panels) and visual judgements (lower panels). Filled (open) circles indicate the mean (and 95% CI) for those participants with the order constant‐variable (variable‐constant), dashed and dotted lines show the respective individual data. Right panel: Proportional biases reflecting the overall integration effect.

To compare these biases between conditions, we collapsed the data across directions of rotation in the form of proportional biases (Fig. 2, right panel). For hand judgements this proportional bias was significantly smaller for condition *variable* (mean and 95% CI: 0.31 [0.21 0.40]) compared to condition *constant* (0.44 [0.34 0.55]; one‐sided paired‐samples t‐test: *t*(27) = 1.92, *p* = 0.032, d = 0.36, B_1_ = 1.9). Similar, for visual judgements the proportional bias was significantly different between conditions *variable* (0.07 [‐0.04 0.19]) and *constant* (‐0.19 [‐0.24 0.01]; *t*(27) = 3.21, *p* = 0.002, d = 0.61, B_1_ = 23.1; for this one‐sided test the alternative statistical hypothesis posited a stronger negative proportional bias and thus a smaller rather than larger mean in condition *constant*). Since the order of conditions was counterbalanced across participants we verified that the difference between conditions was not substantially affected by their order: an ANOVA returned no significant main effect of order (*F*(1,26) = 0.98, *p* = 0.33 and *F*(1,26) = 1.01, *p* = 0.32 for hand and visual judgements respectively) and no interaction between order and conditions (*F*(1,26) = 0.35, *p* = 0.56 and *F*(1,26) = 0.54, *p* = 0.47 for hand and visual judgements respectively).

The mutual attraction of the judged positions of the hand and the visual feedback is an indicator of the overall strength of multisensory integration (Debats et al., 2017b). We estimated this as the difference of proportional biases for hand judgements minus visual judgements, which reflects the fact that mutual attraction results in positive biases for the hand judgements and negative biases for the visual judgements. This overall measure of integration differed significantly between conditions (*t*(27) = 3.57, *p* < .001, d = 0.68, B_1_ = 51.2; individual means and CI’s: *constant* 0.56 [0.38 0.75], *variable* 0.24 [0.09 0.38]) and the Bayes factor provided strong evidence in favour of our main hypothesis that reducing the correlation between hand and cursor trajectories reduces the integration bias.

### Sensory recalibration

We quantified the recalibration biases based on judgements in the unimodal trials of the pre‐ and post‐tests. Figure 3 shows these separately for each condition and for participants with visuo‐motor rotations of +30° and ‐30° in the exposure phase. As compared to the pre‐test, the errors for hand judgements became more positive (negative) in the post‐test after +30° (‐30°) exposure, indicating proprioceptive recalibration. The errors for visual judgements were small.

**Figure 3.**
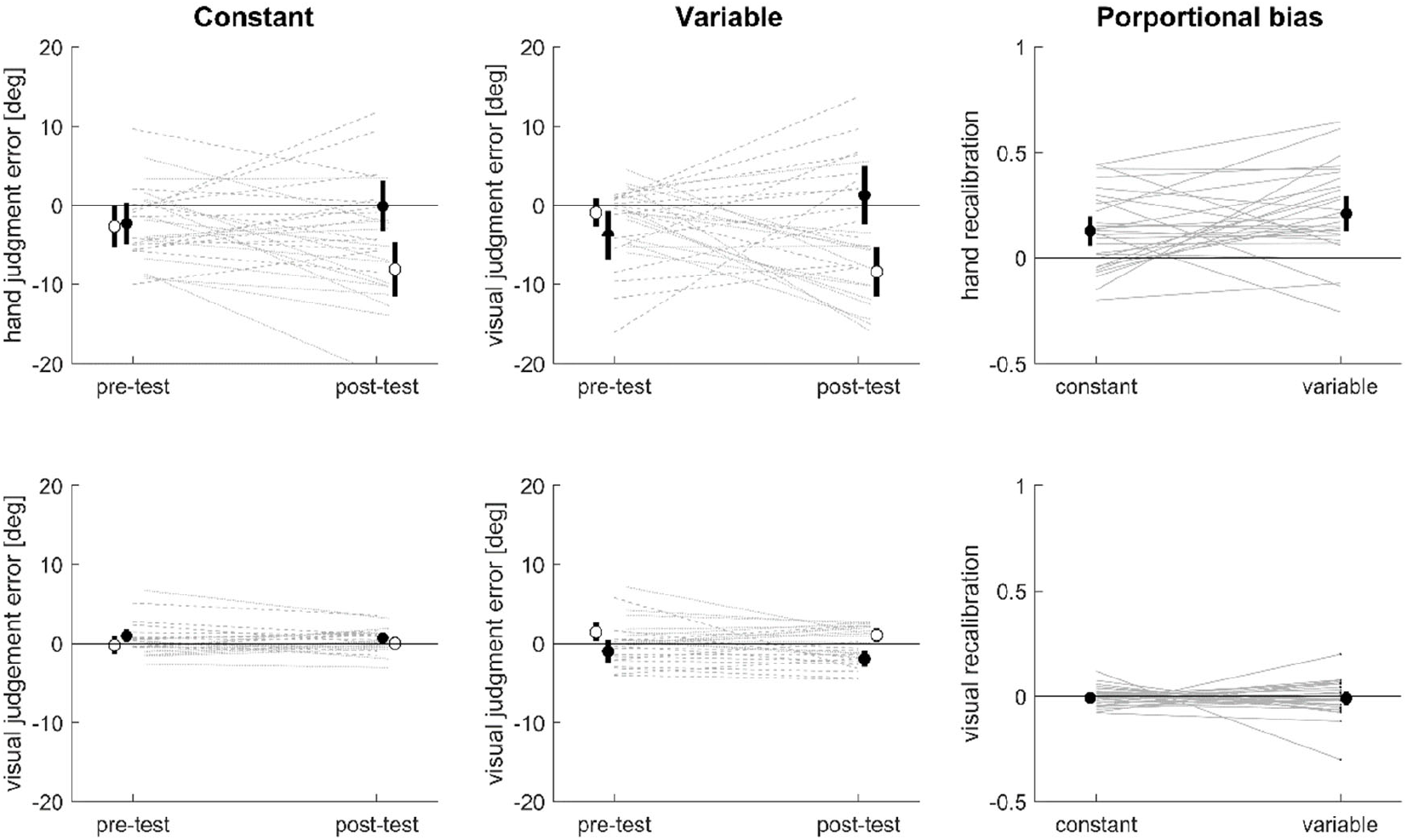
Sensory recalibration. Left and middle panels: Judgement errors in unimodal pre‐test and post‐test trials for hand (upper panels) and visual judgements (lower panels). Filled (open) circles show the means (with 95% CI) for those participants who had been exposed to a visuo‐motor rotation of +30° (‐30°), dashed and dotted lines show the respective individual data. Right panel: Proportional biases reflecting the overall recalibration effect; these are the differences between post‐test and pre‐test, multiplied by ‐1 for exposure to ‐30° and divided by the 30° rotation during the exposure phase.

To summarize these data, we computed proportional biases as the differences between post‐test and pre‐test judgements divided by the visuo‐motor rotation (for ‐30° exposure the differences were multiplied by ‐1 so that positive biases indicate the expected proprioceptive recalibration and negative biases the expected visual recalibration). For hand judgements the proportional biases did not differ significantly between conditions *variable* (mean and CI: 0.21 [0.13 0.29]) and *constant* (0.13 [0.05 0.20]), and the Bayes factor provided ‘strong evidence’ against an attenuated recalibration in condition *variable* (one‐sided paired‐samples t‐test: *t*(27) = ‐1.76, *p* = 0.96, d = ‐0.33, B_0_ = 12.4). Similar for visual judgements the proportional bias was not significantly different between conditions *variable* (‐0.01 [‐ 0.04 0.02]) and *constant* (‐0.01 [‐0.03 0.01]), and the Bayes factor provided moderate evidence against an attenuated recalibration in condition *variable* (*t*(27) = ‐0.07, *p* = 0.53, d = ‐0.01, B_0_ = 5.2). As for integration we verified that the difference between conditions was not substantially affected by their order during the experiment: an ANOVA returned no significant main effects of order (*F*(1,26) = 0.516, *p* = .479 and *F*(1,26) = 0.825, *p* = .372 for hand and visual judgements respectively) and no interaction between order and condition (*F*(1,26) = 1.543, *p* = .225 and *F*(1,26) = 0.688, *p* = .414).

We estimated the overall recalibration strength as the difference of the proportional biases of hand judgements minus the visual judgements. This overall measure of recalibration provided strong evidence against an attenuated recalibration after exposure to degraded local kinematic correlations (t‐test: *t*(27) = ‐1.47, *p* = 0.92, d = ‐0.28 B_0_ = 11.2; individual means and CI’s: *constant* 0.135 [0.061 0.209], *variable* 0.219 [0.128 0.311]), supporting our hypothesis that the within‐trial decorrelation of visual and proprioceptive movements should not affect recalibration.

### Motor adaptation

We quantified motor adaptation based on endpoint deviations in the unimodal trials of the pre‐ and post‐tests. Figure 4 shows these separately for each condition and for participants with visuo‐motor rotations of +30° and ‐30° in the exposure phase. As compared to the pre‐test, endpoint deviations became more negative in the post‐test after +30° exposure and more positive after ‐30° exposure, indicating a negative aftereffect which constitutes motor adaptation.

**Figure 4.**
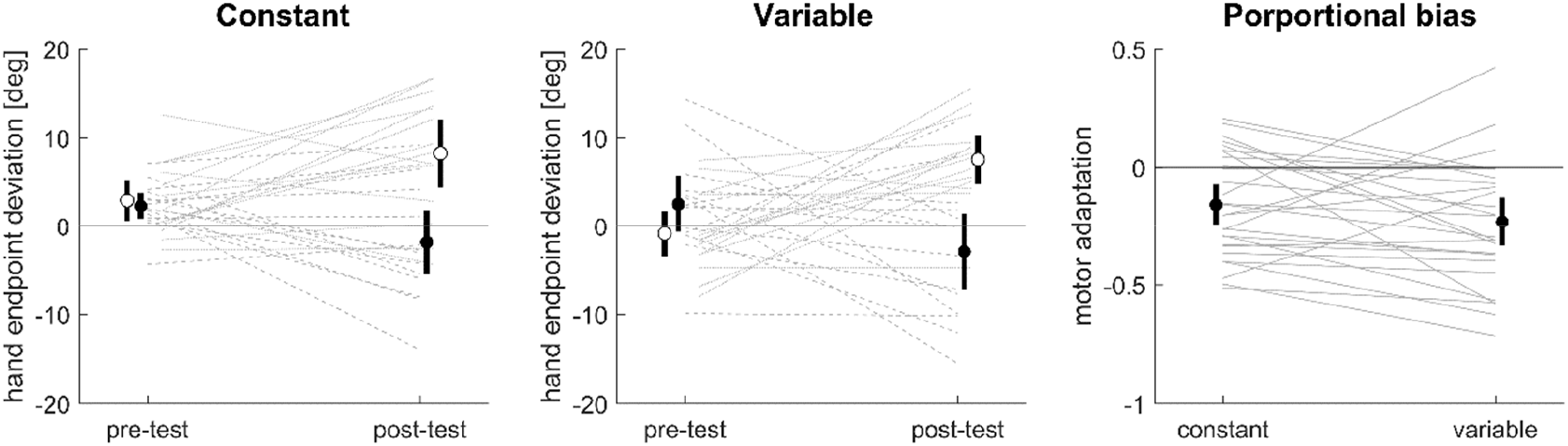
Motor adaptation. Left and middle panels: Endpoint deviations in unimodal pre‐test and post‐test trials. Filled (open) circles show the means (with 95% CI) for those participants who had been exposed to a visuo‐motor rotation of +30° (‐30°), dashed and dotted lines show the respective individual data. Right panel: Proportional endpoint biases reflecting motor adaptation: these are the differences between post‐test and pre‐test, multiplied by ‐1 for exposure to ‐30° and divided by the 30° rotation during the exposure phase. Note that these endpoint biases are negative aftereffects.

The proportional biases for motor adaptation were computed as the differences between post‐test and pre‐test divided by the visuo‐motor rotation of 30° (for ‐30° exposure the differences were multiplied by ‐1 so that negative biases indicate the expected motor adaptation). Numerically the proportional motor adaptation was smaller in condition *constant* (mean and CI: ‐0.158 [‐0.243 ‐0.073]) compared to condition *variable* (‐0.232 [‐0.334 ‐0.129]), which is opposite to what one would expect if degraded kinematic correlations attenuated motor adaptation. A statistical comparison provided ‘strong evidence’ against such an attenuation (*t*(27) = ‐1.51, *p* = 0.93, d = ‐0.26, B_0_ = 11.4). Again this finding was not affected by the order of conditions during the experiment (main effect of order: *F*(1,26) = 0.08, *p* = 0.77, interaction with conditions: *F*(1,26) = 0.20, *p* = 0.65).

### Multisensory integration is affected by exposure to visuo‐motor rotations

In a previous study using a very similar paradigm we observed that the integration bias was abolished after extended exposure to a visuo‐motor rotation of 30° or 0° (Rand and Heuer, 2020). We hence sought to validate this finding in the present data by comparing the total proportional integration bias between pre‐ and post‐tests. Numerically these biases were stronger in the pre‐tests (mean and CI for *constant*: 0.56 [0.38 0.75], for *variable*: 0.24 [0.09 0.38]) compared to post‐tests (0.24 [0.05 0.42] and 0.13 [‐0.04 0.30]). For the statistical comparison with the alternative hypothesis that integration is stronger in the pre‐tests than in the post‐tests we averaged the biases in conditions *constant* and *variable*. The paired‐samples t‐test was significant (*t*(27) = 2.684, *p* = .012, d = 0.51, B_1_ = 7.70), substantiating the finding of reduced multisensory integration after exposure to a series of trials with the same visuo‐motor rotation of feedback‐endpoints relative to hand‐movement endpoints.

## Discussion

Using a visuo‐motor task as model of multisensory perception we probed how reducing the spatio‐ temporal correlation of the information received by two senses affects multisensory integration, recalibration and also motor adaptation. By reducing the kinematic correlation of the felt hand movement and the seen trajectory of the visual feedback we reduced the moment‐by‐moment congruency between visual and proprioceptive signals in a visuo‐motor task, but left intact their correspondence at the start and end of each movement. We expected this manipulation to reduce integration but not recalibration and motor adaptation. Our data confirmed this hypothesis. In an ancillary analysis, we found that prolonged exposure to visuo‐motor information with a constant offset of movement endpoints reduces the integration bias, an observation also made in a previous study (Rand and Heuer, 2020).

### Effects of kinematic (de‐)correlation on integration and recalibration

Multisensory integration supposedly serves to reduce the discrepancy between two redundant sensory signals received more or less at the same time and pertaining to a common cause in our environment (Kording et al., 2007; Rohe and Noppeney, 2015b, a; Odegaard et al., 2017; Noppeney, 2021; Shams and Beierholm, 2022). This predicts that manipulations reducing the momentary evidence for two signals to arise from the same cause should reduce integration (Debats et al., 2017a; Debats et al., 2023). In contrast, multisensory recalibration is a process that supposedly serves to reduce persistent or constant discrepancies between sensory estimates (Recanzone, 1998; Bruns, 2019; Noppeney, 2021; Bruns et al., 2022). Though recalibration emerges on multiple time scales, the bias observed following prolonged exposure is the most well‐studied and robust effect and the one investigated here. This recalibration bias is likely driven by estimates of sensory contingencies on longer time scales and may not be affected by manipulations on short time scales (Bosen et al., 2017; Bosen et al., 2018).

To directly test this prediction, we implemented a manipulation of the kinematic correlation of proprioceptive and visual signals during individual movements that reduced their moment‐by‐moment correlation during each individual trial. In contrast, the more global or long‐term correlation between the end positions of the trajectories of hand and feedback, computed across trials, remained perfect. In support of our hypothesis we found that the integration bias was reduced by the manipulation of kinematic correlations, while recalibration was not. Hence our data support the conclusion that integration and recalibration are shaped by evidence for a common cause of two signals on different time scales. Our data also suggest that motor adaption is similarly robust to reduced kinematic correlations as is multisensory recalibration and hence is guided by the sensory contingencies on possibly the same time scales.

These results promote a number of additional hypotheses, some of which have already been tested in previous studies, and some which could be tested in the future. In the present experiment we manipulated the kinematic correlation on a short time scale but kept those on the longer time scale intact. The opposite manipulation, keeping the within‐trial correlation of the two signals intact but manipulating that on longer (trial by trial) time scales should produce the opposite results as obtained here; that is, should preserve integration. Such a condition has been studied in the visuo‐motor paradigm by (Rand and Heuer, 2019a), who tested integration for individual visuo‐motor rotations after a series of trials featuring the same rotation, that is, after the experience of a perfect trial‐by‐trial correlation of proprioceptive and visual feedback. In this condition the integration bias was equivalent to that in a standard condition with variable visuo‐motor rotations between trials, suggesting that indeed variations of longer term common‐cause evidence do not affect multisensory integration.

In contrast, varying the long‐term correlation between senses should modulate recalibration and motor adaptation. For motor adaptation relevant data are available. Indeed, across variations of the overall experimental design the adaptation effect seems to be attenuated when visuo‐motor rotations are variable rather than constant (Fernandes et al., 2012; Saijo and Gomi, 2012; Butcher et al., 2017; Albert et al., 2021). According to our hypothesis, this reduction is a consequence of reduced common‐cause evidence and thus a reduced binding of the bimodal position estimates. Reduced binding again could result in the reduced error sensitivity that has been shown for variable visuo‐motor rotations (Avraham et al., 2020; Albert et al., 2021) – after all there should be little benefit from adjusting movement directions as a consequence of visually indicated errors that depend on random spatial offsets between hand and cursor positions. Similarly detailed analyses of sensory recalibration are missing, but there is evidence that variations of motor adaptation often imply concomitant variations of recalibration (Tsay et al., 2022).

The moment by moment evidence for a common cause can be manipulated based on the kinematic correlation as done here, but also by replacing the continuous visual feedback by terminal feedback that is presented only at the end of each outward movement. Consistent with the present hypothesis, integration is attenuated with terminal compared to continuous visual feedback (Debats and Heuer, 2018), while recalibration is largely unaffected (Barkley et al., 2014; Ruttle et al., 2022). For motor adaptation the differences between continuous and terminal visual feedback vary between studies: while in some experiments adaptation was reduced for terminal feedback (Hinder et al., 2008; Taylor et al., 2014), it was enhanced in others (Bernier et al., 2005; Heuer and Hegele, 2008). The reasons for the divergent findings remain to be investigated.

### Multisensory integration is attenuated after exposure to constant visuo‐motor rotations

Multisensory integration is known to be sensitive to the overall experimental context in which it is probed, and the bias observed in particular trials can depend on the nature of preceding trials in an experiment (Gau and Noppeney, 2016; Rand and Heuer, 2018; Rohe et al., 2019). For example, with a larger range of intermodal discrepancies we observed weaker integration in the visuo‐motor task (Debats et al., 2023), but stronger integration in an audio‐visual ventriloquism paradigm (Park and Kayser, 2022). In line with such a context dependency of integration we have previously observed that prolonged exposure to constant discrepancies between hand movements and visual feedback reduces the integration bias (Rand and Heuer, 2020). In that study, the multisensory integration observed in test trials using variable rotations was reduced following prolonged exposure to trials featuring a constant visuo‐motor rotation. The data obtained in the present experiment confirm this result.

Such context sensitivity in the visuo‐motor paradigm can be explained based on the notion that visual consequences of movements are combinations of actual sensory input and prior expectations (Diedrichsen et al., 2010; Shimansky and Rand, 2013). Extended exposure to a constant rotation may strengthen this prior. Hence in test trials with variable rotations the prior referring to a constant rotation may dominate over the variable sensory evidence. At the same time, exposure to constant rotations should also strengthen the evidence of a common cause of visual and proprioceptive signals. The reduced integration in spite of common‐cause evidence shows that integration is not sensitive to the causal evidence accumulated across trials. Rather, the results of the present study suggest that the evidence about a common cause guiding integration is based on the momentary sensory information obtained during individual movements.

### Conclusion

The data obtained here suggest that multisensory integration and sensory recalibration deal with multisensory discrepancies in distinct ways: integration is engaged by variable discrepancies, prompted by local common‐cause evidence, and becomes ineffective when variable sensory discrepancies become dominated by strong prior expectations of specific discrepancies. In contrast, recalibration (and motor adaptation) are engaged by consistent discrepancies, prompted by global common‐cause evidence, and become less effective when variability of intermodal discrepancies prevents the development of strong priors.

## Acknowledgements and funding

This study was funded by the Deutsche Forschungsgemeinschaft (DFG KA2661/2‐1). The authors declare that they have no conflict of interests. We would like to thank Lisa Stetza and Jenny Jakisch for help with data acquisition.

## Data availability

The data and relevant Matlab code for producing the figures will be made available under https://github.com/christophckayser

